# Genomic Insights into *Wolbachia* Strain wCin2USA1 Reveal Promising Cytoplasmic Incompatibility Potential and Next-Generation Dengue Biocontrol

**DOI:** 10.1101/2025.08.22.671735

**Authors:** Istiaque Zaeem, Nurnabi Azad Jewel, Mohimenul Haque Rolin, Daniyal Karim, Arzuba Akter, Shakhinur Islam Mondal

## Abstract

Dengue fever poses a growing public health challenge globally, resulting in significant morbidity and economic burden. The use of *Wolbachia*-mediated biocontrol represents a promising, cost-effective, and environmentally sustainable strategy for managing dengue transmission. However, the susceptibility of existing *Wolbachia* strains utilized in controlling *Aedes aegypti* necessitates the investigation of novel strains to enhance dengue control efficacy. This study aimed to identify potential alternative *Wolbachia* strains for dengue control by comparing the genomes of seven *Wolbachia pipientis* strains: wMel, wAlbB, wlrr, wHm-C, wAnm, ant7, and a strain isolated from *Aedes aegypti*. We conducted comprehensive genomic analyses, including phylogenetic assessments, metabolic pathway evaluations, and characterization of the Cytoplasmic Incompatibility Factor (*Cif*) genes. Our analyses identified the strain wCin2USA1 as a strong candidate for alternative dengue control strategies. This strain demonstrated remarkable genomic similarities to wMel, an already established strain used as biocontrol for *Aedes aegypti.* Importantly, this strain presented two distinct pairs of Cif genes from different monophyletic types, each homologous to the Cif genes found in wMel and wAlbB. This genetic architecture suggests a high degree of compatibility and demonstrates promising potential for the suppression of *Aedes aegypti* populations through the induction of Cytoplasmic Incompatibility. The presence of multiple intact prophage regions also suggests greater adaptability compared to established strains. Our findings support the hypothesis that wCin2USA1 could serve as an effective biocontrol agent against dengue transmission. This work provides critical insights into developing innovative *Wolbachia*-based interventions aimed at mitigating the persistent threat posed by dengue fever. Future research should concentrate on optimizing release methodologies, evaluating ecological impacts, and assessing the strain’s effectiveness against the dengue virus.

**Author Summary:** Dengue fever is one of the fastest-growing mosquito-borne diseases in the world, causing illness and economic challenges in many countries. Current mosquito control methods, such as insecticides, are often costly, less effective over time, and harmful to the environment. An alternative approach uses naturally occurring bacteria called Wolbachia, which live inside insects. When certain Wolbachia strains are introduced into mosquitoes, they can reduce the insects’ ability to spread viruses like dengue. In our study, we compared the genetic makeup of several Wolbachia strains to identify new candidates that may work better for controlling dengue. We discovered that a strain called wCin2USA1 has strong potential because it shares important features with two strains already used successfully, while also having unique advantages. These include genes that help prevent mosquitoes from reproducing normally when carrying different Wolbachia strains, which can reduce mosquito populations. Our findings suggest that wCin2USA1 could be developed as a new, environmentally friendly tool to help reduce dengue transmission.

## Introduction

Dengue fever is a highly contagious viral disease transmitted by vectors, especially by *Aedes aegypti*. The rapid rise of dengue poses a significant global public health challenge, as nearly half of the world’s population is now at risk, with an estimated 100 to 400 million infections occurring annually (WHO, 2024). The adoption of numerous countermeasures has been undertaken to control the spread of dengue fever. Present management endeavors concentrate on decreasing populations of the dengue vector using a variety of biocontrol methods. A major biocontrol strategy includes the introduction of endosymbiotic bacteria that disrupts various biological functions within the hosts’ bodies, including dysregulating their sexual cycles by inducing Cytoplasmic Incompatibility (CI) as well as reducing their ability to transmit illnesses, and impeding the growth stages of the vector species (Wilke & Marrelli, 2015). Consequently, these alterations lead to suppression of the overall vector population. Notably, one particularly impactful group of bacterial agents employed in this context is *Wolbachia*. *Wolbachia* is a genus of Gram-negative bacteria that infects a wide range of arthropod species, typically functioning as an intracellular parasite, though it also exhibits mutualistic relationships in filarial nematodes. It is one of the most prevalent endosymbiotic microbes in arthropods and is recognized as one of the most widespread reproductive manipulators in the biosphere (Werren et al., 2008). *Wolbachia* has complex interactions with its hosts, with some host species unable to reproduce or survive without its colonization. Cytoplasmic Incompatibility (CI) caused by *Wolbachia* is a reproductive barrier where uninfected female *Aedes aegypti* cannot produce viable offspring when they mate with male carriers of *Wolbachia*. Although there are no verified cases of *Wolbachia* being transmitted vertically purely through males, certain unidentified modifications lead to fatal consequences during the early stages of embryo development after fertilization (Sicard et al., 2019). Infected female mosquitoes can produce healthy offspring whether they mate with *Wolbachia*-carrying or uninfected mates. Infected individuals harboring *Wolbachia* experience improved survival chances, resulting in a selection advantage that leads to a higher prevalence of the infection in affected groups (Joshi et al., 2014).

Genetic studies have identified two genes, CifA and CifB, associated with Cytoplasmic Incompatibility (CI) in *Wolbachia*. These genes are frequently located together in prophage eukaryotic association modules (EAMs) and are thought to disseminate through horizontal transmission among *Wolbachia* strains because they exist in prophage regions (Kaur et al., 2022). There are two main models explaining CI: the toxin-antidote (TA) model and the host-modification (HM) model. The TA model suggests that CifB alters paternal chromatin in embryos, countered by maternally inherited CifA, leading to incompatibility between *Wolbachia* strains with different Cif homologues (Beckmann et al., 2019). The HM model proposes that CifB modifications occur during spermatogenesis, with CifA as a co-factor, suggesting rescue without Cif peptide packaging into sperm. This model lacks a clear explanation for the diverse compatibility patterns seen in nature (Shropshire et al., 2020).

While the currently established strains, wMel and wAlbB primarily, continue to be used as biocontrol agents to control mosquito populations, their efficiency and stability over long-term is becoming questionable. It is also possible that escape mutations will emerge in dengue or other arboviruses, which will reduce the efficiency of transmission blocking by current *Wolbachia* strains (McNamara et al., 2024). In this context, it is essential to continue research and extend our understanding about other plausible strains to counter these issues.

According to available studies, wMel is generally more effective in blocking dengue virus replication than wAlbB, although both strains have been reported to decrease the transmission potential of dengue virus (Xue et al., 2018). Additionally, wMel does not appear to impose a substantial fitness cost on the mosquitoes, whereas wAlbB has been associated with slightly lower fitness benefits compared to wMel (Sadanandane et al., 2022). However, both strains have proven successful in reducing dengue virus transmission when released in the field.

A concerning fact is number of recent studies have demonstrated that CI, which is an essential quality for the efficacy of wMel, is diminished by the cyclical heat stress that can be a characteristic of field environment (Ross, Ritchie, Axford, & Hoffmann, 2019). Once a threshold of 35℃ is achieved, high temperatures cause the infection to be less efficient and limit the hatch rate of eggs that are infected with wMel. Additionally, high temperatures reduce the density of wMel in adults, which hinders the transfer of the *Wolbachia* from parent to offspring (Ross et al., 2019). However, the other strain, wAlbB, has been successfully established in Malaysian populations of *Aedes aegypti*. It is shown to be less susceptible to high rearing temperatures and has been successfully established in Malaysian populations (Nazni et al., 2019).

In events where global warming continues to pose a constant threat, the proportion of the world that is exposed to diseases transmitted by mosquitoes that are fatal, increase because of rising temperatures. Moreover, it may also reduce the usefulness of the *Wolbachia*-mediated biocontrol by the wMel strain that has been shown to be effective in controlling vector population (Ross et al., 2019). Considering all these factors, it is not irrational to say that it is high time we started looking for more effective strains of *Wolbachia* for the reassurance that *Wolbachia*-mediated biocontrol continues to be a resilient technology against mosquito-borne diseases in the face of near-term climate change.

Here, we examine the genomes of eight strains of *Wolbachia pipientis* for comparative analysis of genetic characteristics and features of Cytoplasmic incompatibility factor genes or Cif genes that are mainly linked to the capacity to induce cytoplasmic incompatibility (CI) to find more effective alternate strains alongside wMel and wAlbB. Comparative analyses of the *Wolbachia* genomes have been undertaken to distinguish their distinctive genetic attributes, whereas assessments of Cif genes seek to illuminate characterization of Cif genes instigated by differing *Wolbachia* strains responsible for disrupting host reproduction. We show that a particular strain, wCin2USA1 showcases highly similar genomic properties with wMel strain yet possesses distinctive sets of functional Cif genes. Additionally, our study depicts that wCin2USA1 has multiple intact prophage regions, implying its versatility and adaptability to different environments. Potentially, wCin2USA1 strain can be one of the answers to the ever-threatening mosquito-borne diseases in the upcoming future whereas currently established strains may substantially be ineffective over time.

## Materials and Methods

### Retrieval and Curation of *Wolbachia* genomic sequences

A total of 26 *Wolbachia pipientis* whole-genome sequences were retrieved from the NCBI database. To reduce redundancy within this dataset, CD-HIT (Fu et al., 2012) was employed to cluster and remove highly similar sequences. A stringent sequence identity threshold of 99% (-c 0.99) was applied to ensure the exclusion of near-identical genomes.

### Genome Annotation

Total eight *Wolbachia* genomes including: wMel, wAlbB, wAnm, wlrr, ant7, wHm-c and *Aedes aegypti* isolate strains were identified as suitable for analysis, underwent various annotation processes to ensure uniformity and accuracy in functional annotation. PROKKA (Seemann, 2014) was utilized via the command-line interface to for functional annotation.

### Genome Mapping

To generate a BLAST comparison image among all eight strains, showcasing their similarities Blast Ring Image Generator (BRIG) (Totsika et al., 2011) was utilized. For the visualization of genome conservation among all strains, EasyFig (Sullivan et al., 2011) was used. FastANI v1.34 (Jain et al., 2018) was utilized to visualize pairwise genome comparison between *Aedes aegypti* isolate strain and other relative strains to associate genome conservation among them.

### Phylogenetic Tree Construction and Analysis

To elucidate the evolutionary relationships among *Wolbachia* strains and closely related bacteria, orthologous and paralogous gene families were assigned for 8 *Wolbachia* strains and 20 outgroup strains from the class *Alphaproteobacteria* that are closely related to *Wolbachia* using OrthoFinder v2.5.5 (Emms & Kelly, 2015). The orthologous groups containing only one gene in each strain or single-copy genes were selected to construct the phylogenetic tree. The protein sequences of each orthologous group were independently aligned using MAFFT v7.520 (Rozewicki et al., 2019). The phylogenetic tree was constructed based on maximum likelihood (ML) using with the resultant alignment used as input into iq-tree v2.0.6 (Nguyen et al., 2015) using default parameters. The tree was visualized using iTOL v6 server (https://itol.embl.de).

### Orthologous Gene Cluster Analysis

To identify and visualize orthologous gene clusters among the selected genomes, OrthoVenn3 (https://orthovenn3.bioinfotoolkits.net/home) was used. The protein sequence files (in FASTA format) of the relevant bacterial strains were uploaded to the platform. The default parameters were used: E-value cutoff of 1e-5 and an inflation value of 1.5 for MCL clustering.

The resulting orthologous clusters were then visualized using the interactive Venn diagram provided by the platform. The output figure was exported directly from OrthoVenn3 and used to illustrate comparative genomic relationships in the final results.

### Metabolic Pathway Construction and COG Analysis

To comprehensively analyze the metabolic pathways, all the anticipated protein-encoding genes from these strains were submitted to the EggNOGmapper (v2.1.12) (Huerta-Cepas et al., 2017) and the KEGG Automatic Annotation Server (KAAS) (Moriya et al., 2007) for automated functional annotation. The existence and the absence of genes within these pathways were manually evaluated and displayed as a heatmap.

### Cytoplasmic Incompatibility Factor (Cif) Genes Characterization

The nucleotide sequences for the CI factor (Cif) A and B genes for all five monophyletic types of *Wolbachia* strains were retrieved from the study conducted by Martinez et al. (Martinez et al., 2019). Based on high confidence level and accuracy (probability score > 99%, e-value < 1E-10, coverage = 100), Cif gene homologues in the eight Wolbachia strains analyzed in this study were identified.

Partially sequenced *cif* genes were excluded from the analysis. Complete nucleotide sequences of *cifA* and *cifB* genes were used to construct a custom database using DIAMOND (Buchfink, Reuter, & Drost, 2021). The eight *Wolbachia* genomes were then queried against this database using DIAMOND with the --ultra-sensitive parameter to identify *cif* gene homologs.

The nucleotide sequences for the CifA and CifB genes were aligned separately using the program MAFFT. The separate nucleotide sequence alignments were concatenated using SeqKit (Shen et al., 2016). Poorly aligned parts were trimmed using TrimAl. The concatenated alignment was then used as input for into iq-tree v2.0.6 (Nguyen et al., 2015). The ML tree was inferred using IQ-TREE 1.5.5 ran with ModelFinder and tree reconstruction (Nguyen et al., 2015). The outputted Newick formatted tree was then annotated using using iTOL v6 server (https://itol.embl.de).

## Results and Discussion

### *Wolbachia* Strain wcin2USA1 Genomic Composition with Distinctive Features

Our study provided a comprehensive analysis of eight strains of *Wolbachia pipientis* that have been sequenced on a whole genome level. Among *Wolbachia pipientis* strains, wMel and wAlbB are the most commonly used in biocontrol efforts to reduce flavivirus transmission in Aedes aegypti (Sadanandane et al., 2022).

All eight *Wolbachia* strains show almost similar genome sizes ranging from 1.1 Mb to 1.6 Mb. The strain wCin2USA1 along with the strain ant7 showcases a large genome with a high number of gene content, 1545 and 1579 respectively shown in Table 1.

**Table 1.**
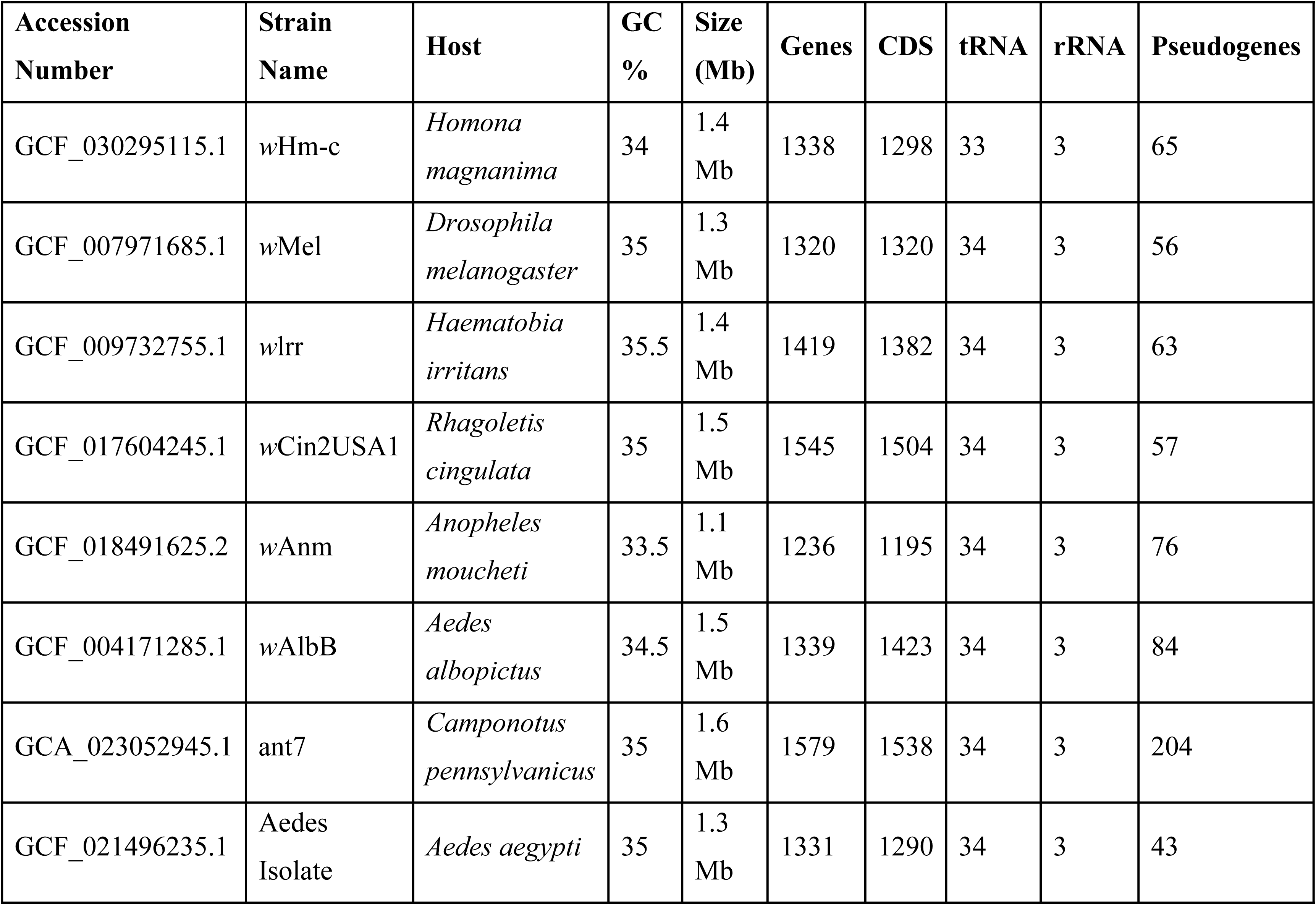
General genomic features including accession numbers, strain names, host organisms, GC content, genome size, number of genes, coding sequences (CDS), transfer RNA (tRNA), ribosomal RNA (rRNA), and pseudogenes of eight distinctive *Wolbachia* strains.

A circular comparative genomic map of the *Wolbachia pipientis* strain isolated from *Aedes aegypti* (1,267,786 bp) aligned against eight reference *Wolbachia* genomes, including wMel, wCin2USA1, wLtr, ant7, wAlbB, wHm-c, and wAnm (Fig 1). Sequence identity levels, depicted through concentric rings, range from 50% to 100%, highlighting regions of conservation and divergence. The inner tracks display GC content and GC skew, with shifts marking the replication origin and terminus.

**Fig 1.**
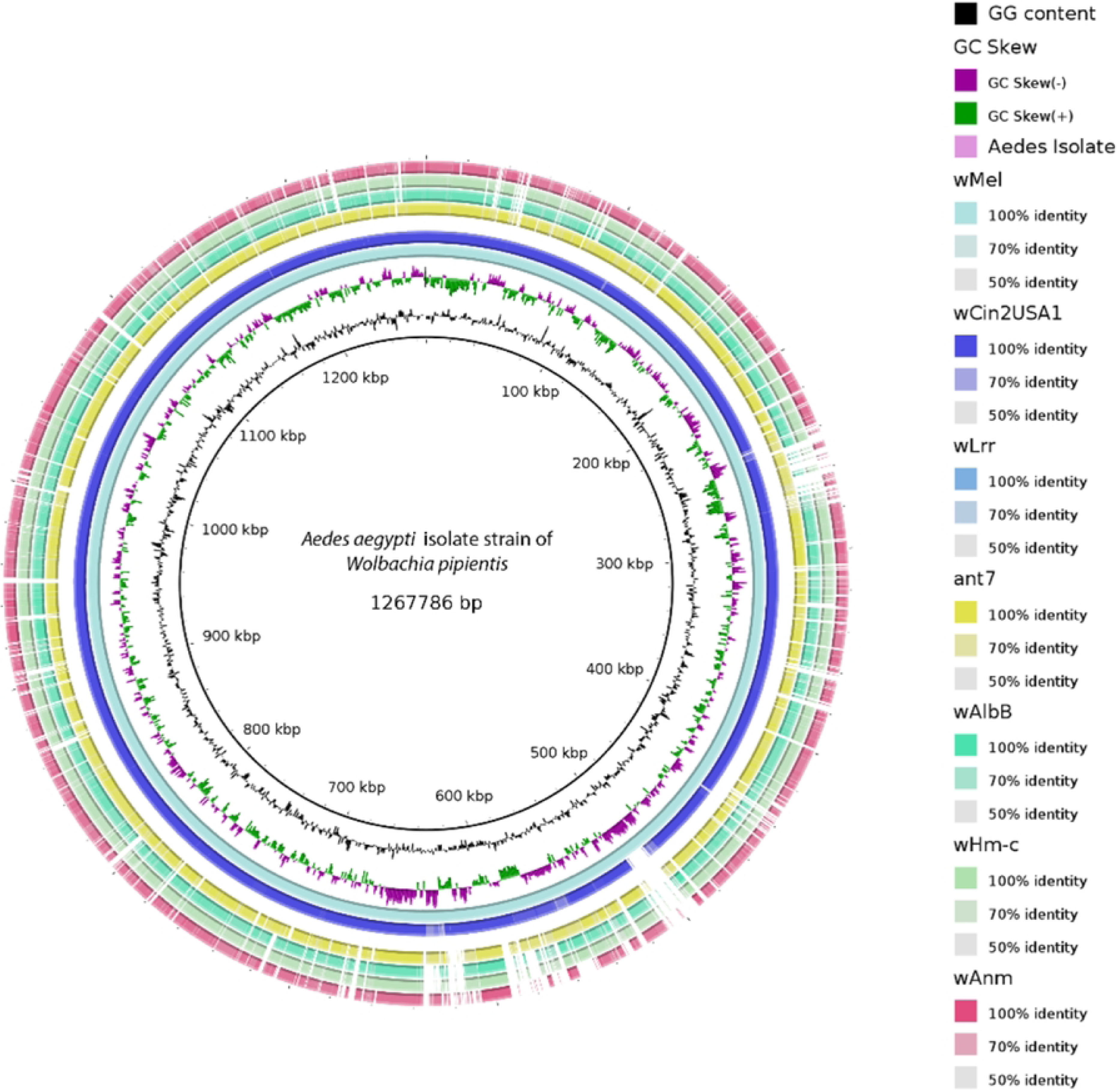
A comparative genomic view of the *Wolbachia* strains based on BLASTn. The innermost and second-innermost circles show the GC content (black) and the GC skew. The GC-skew and GC-content values were calculated using a sliding-window (window size = 1,000 bp; step size = 500 bp).

We can notice a strong genomic conservation between the *Aedes aegypti* isolate and strains wMel, wCin2USA1, suggesting close evolutionary relationships. In contrast, wAnm and ant7 exhibit notable divergence, characterized by fragmented identity blocks, indicating more distant evolutionary relationships or ecological specialization. Strains wHm-c and wLtr show intermediate similarity to the Aedes isolate.

Genome comparison revealed that all *Wolbachia* strains have a very similar GC content percentage, ranging from 33.5% to 35.5%. This narrow range of GC content suggests that GC content among these strains is conserved by purifying selection, and they share a common ancestry (Bohlin et al., 2017).

The wCin2USA1strain showcased an increased gene content when compared to wMel strain. This increased gene content may be due to the acquisition of new genes through horizontal gene transfer (HGT) or the duplication of existing genes, which can lead to functional diversification and adaptation to specific ecological niches.

It is also important to note that larger genomes may contain more non-coding regions or pseudogenes, which can be difficult to attribute to specific functions as it is seen from Table 1. that the ant7 strain possess 204 pseudogenes, which is very high compared to other strains.

The strain wCin2USA1 shows only 57 pseudogenes despite having a large genome and high gene numbers compared to the other strains. This implies that wCin2USA1 can possibly have higher functional specialization and adaptability to different ecological niches than other strains (Pushker et al., 2004).

### Evolutionary Distinctiveness and Closeness of wCin2USA1 to wMel strain

One of the findings of this study is that the *Wolbachia* strains analyzed do not form a monophyletic group within their designated clade, but rather exhibit multiple branching patterns, highlighting considerable genetic divergence among them. Although all eight strains cluster within the same broader clade when placed in a phylogenetic tree alongside other Alphaproteobacteria, their internal branching structure suggests that while they likely descended from a common ancestor, they have undergone substantial evolutionary diversification shown in Fig 2.

**Fig 2.**
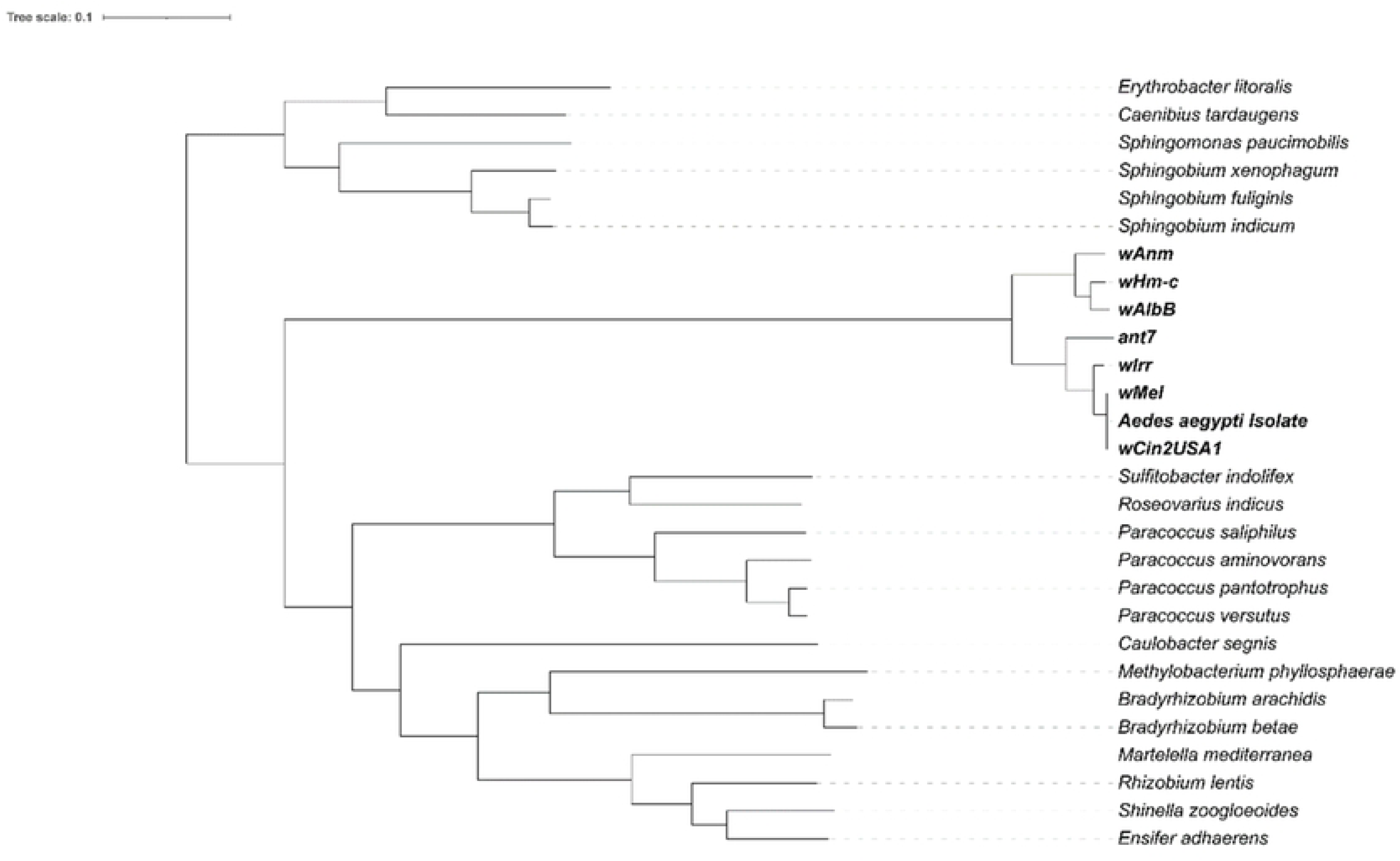
Neighbor-joining maximum-likelihood tree from whole-genome alignments of 28 alphaproteobacterial sequences, including eight *Wolbachia* strains. The tree is midpoint rooted. The scale bar indicates relative branch lengths of the phylogenetic tree. This phylogenetic tree showed that the *Wolbachia* endosymbionts formed a cluster of strong supported monophyletic group where the eight *Wolbachia* strains (bold) are placed in the same clade.

Within the *Wolbachia* grouping, the strains are clearly divided into two distinct subclades. The first subclade comprises ant7, wIrr, wCin2USA1, the Aedes aegypti isolate, and wMel, all of which show relatively short phylogenetic distances indicative of close evolutionary relationships. The second subclade includes wAlbB, wAnm, and wHm-c, which cluster separately, suggesting a divergent evolutionary path within the *Wolbachia* lineage (Fig 3).

**Fig 3.**
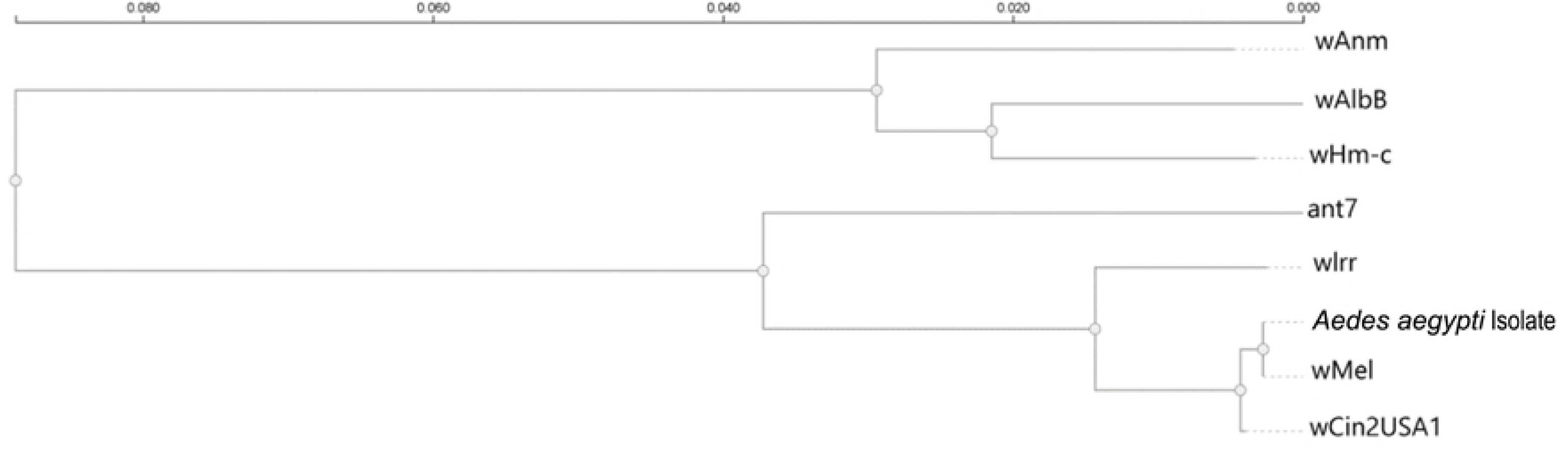
Maximum likelihood phylogenetic tree of eight *Wolbachia* strain. *Aedes aegypti* isolate strain and wMel strain cluster in a sub-clade along with the wCin2USA1 strain. It suggests that the wCin2USA1 may be heavily similar to these strains compared to the other strains. Among the other strains, the wlrr and ant7 strain are slightly distanced from the *Aedes aegypti* isolate strain while the wAlbB and wHm-c strains are in the same cluster together in another sub-clade. The wAnm strain belongs to the furthest branch or sub-group from other strains.

The phylogenetic placement of wCin2USA1 offers valuable insights into the evolutionary diversification and symbiotic potential of *Wolbachia* strains. In our analysis, wCin2USA1 clusters tightly with wMel, a well-characterized strain isolated from *Drosophila melanogaster* and widely used in vector control programs due to its capacity to induce cytoplasmic incompatibility and suppress arboviral replication. This close evolutionary relationship suggests that wCin2USA1, originally isolated from the cherry fruit fly *Rhagoletis cingulata*, may share key genomic features and symbiotic traits with wMel, despite their association with phylogenetically distinct insect hosts.

### Proteomic Divergence Despite High Genomic Identity Between wCin2USA1 and the wMel Strain

Comparative genomic analysis between *Wolbachia pipientis* strain wCin2USA1 and a *Wolbachia* strain wMel revealed a high degree of nucleotide conservation, with an Average Nucleotide Identity (ANI) of 98% (Fig 4).

**Fig 4.**
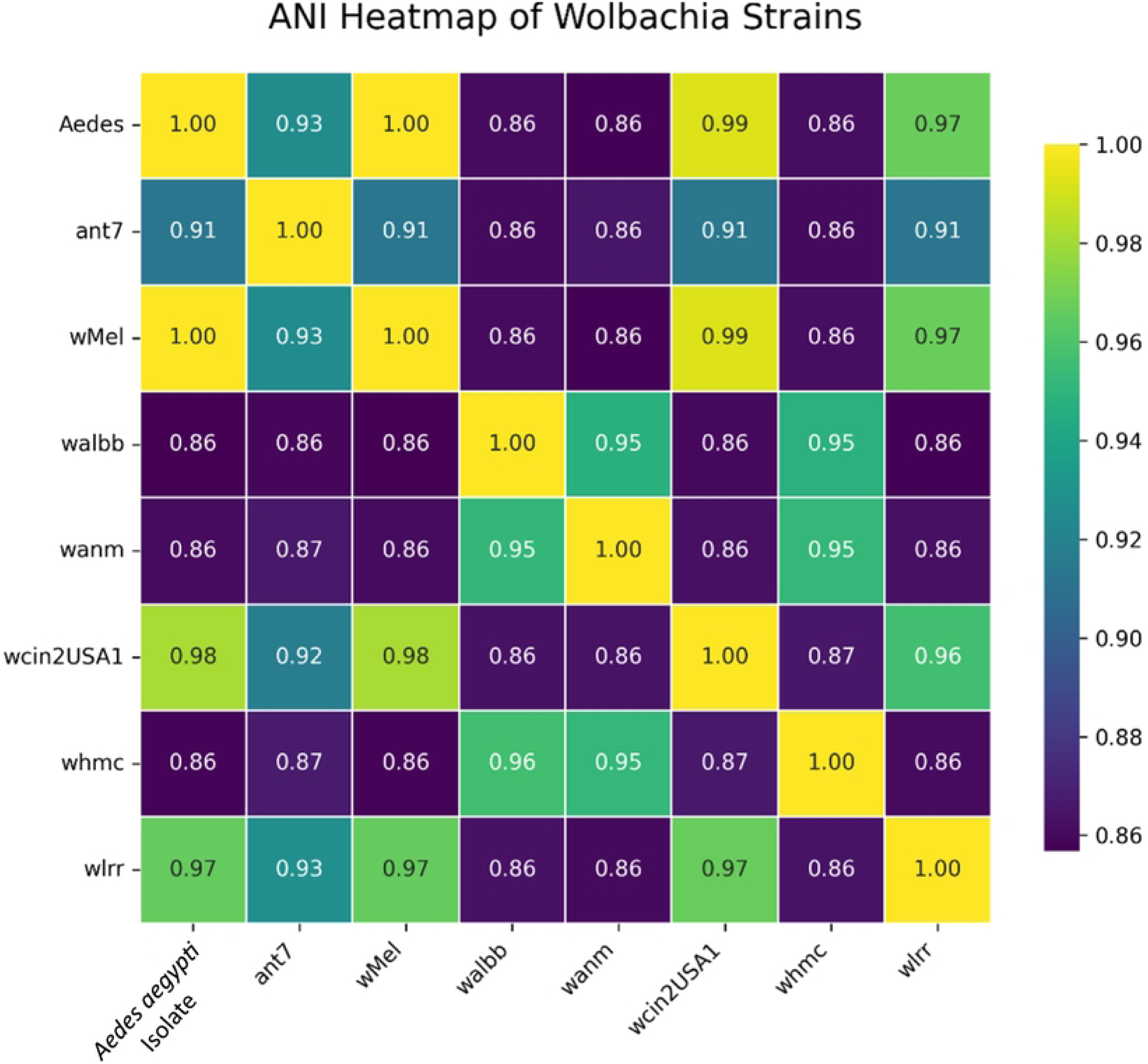
Heat map of average nucleotide identity (ANI) based on whole genome sequence of 8 strains of *Wolbachia*. ANI values (%) are shown in the matrix and these values are indicated by the color intensity ranging from 1 to 0.86.

Synteny analysis further demonstrated extensive collinearity, with large, conserved blocks spanning the majority of both genomes (Fig 5). However, multiple genomic inversions and rearrangements were also observed, indicated by crossing ribbons and shifted syntenic blocks in the alignment. These structural variations, while not extensive, highlight genomic plasticity that could influence gene regulation and expression profiles (Merrikh & Merrikh, 2018).

**Fig 5.**
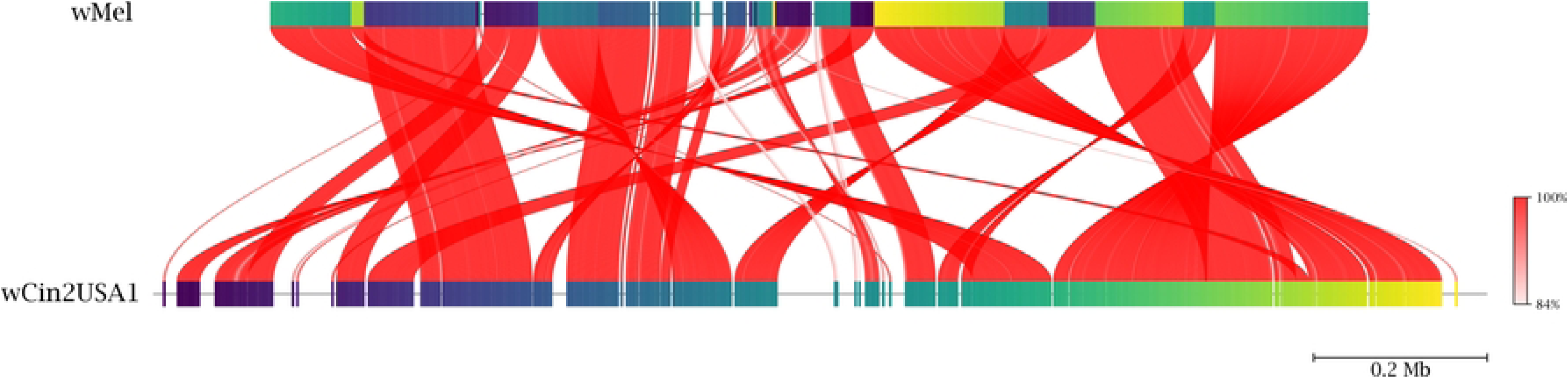
Synteny plot depicting genomic alignment between *Wolbachia pipientis* strain wCin2USA1 (bottom) and a *Wolbachia* strain wMel (top). Red ribbons denote homologous regions, with color intensity indicating sequence identity (84%– 100%). The high degree of synteny and sequence conservation underscores the close genomic relationship between the two strains.

Despite the high ANI similarity, these structural differences may translate into divergent proteomic profiles. Such rearrangements can impact key phenotypic traits, including host-microbe interactions and cytoplasmic incompatibility potential. While wCin2USA1 is highly similar to the wMel strain at the nucleotide level, it likely exhibits functionally distinct characteristics at the proteomic level.

A total of 748 orthologous gene clusters were identified as conserved across all analyzed Wolbachia strains, representing the species’ core genome (Fig 6). Analysis using an UpSet plot revealed that the strains wMel, wCin2USA1, wLrr, and the Aedes aegypti isolate collectively shared 72 orthologous clusters, the highest among all strain combinations. Notably, 47 orthologous clusters were exclusively shared among wMel, wCin2USA1, and the *Aedes aegypti* isolate, representing a significantly enriched subset. Frequent co-occurrence of wMel and wCin2USA1 within shared gene clusters underscores a high degree of genomic similarity between these two strains, suggesting a closer evolutionary relationship compared to other Wolbachia lineages examined.

**Fig 6.**
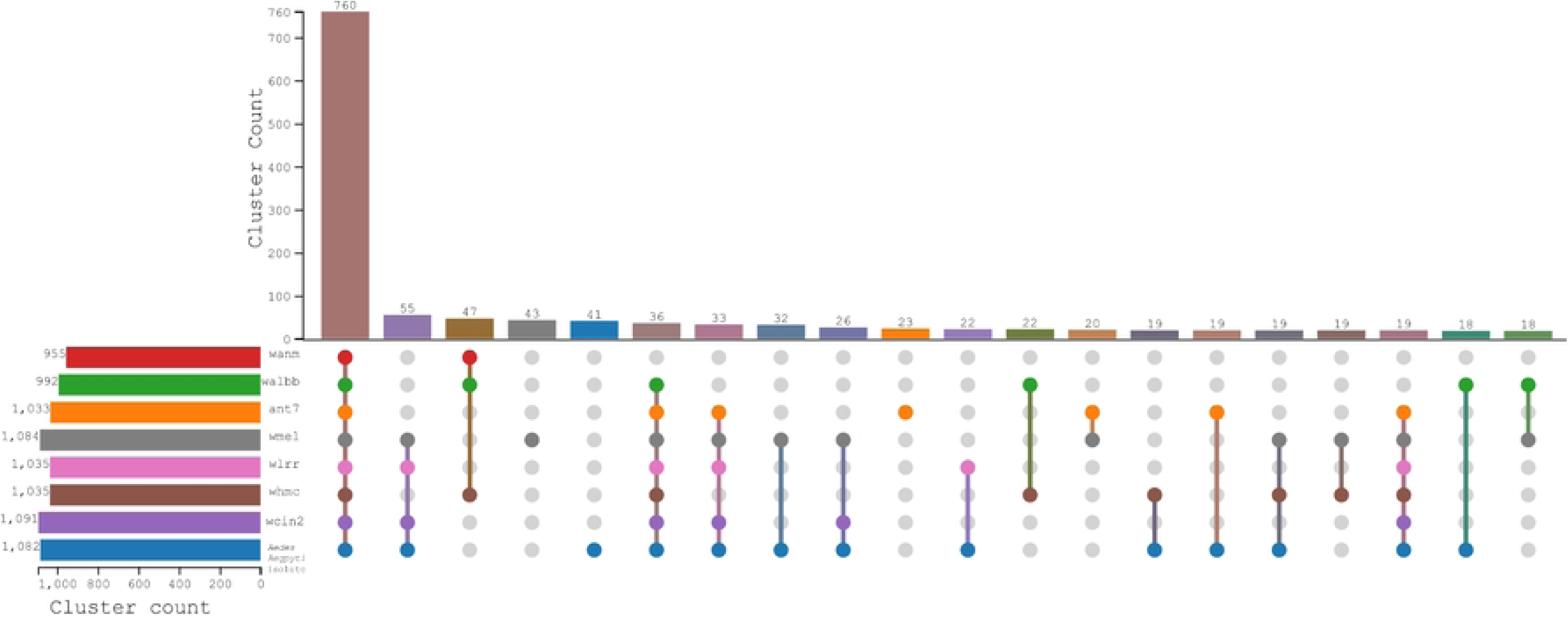
Overview of identified orthogroups amongst *Wolbachia* shown via UpSet plot. Each genome (one per row at the bottom half of the figure:) is treated as a set containing a certain number of orthogroups (denoted by the bar graph on the bottom left of the image). *Wolbachia* genomes are colour-coded based on their strain. An intersection of all 8 *Wolbachia* genomes was identified as containing the vast majority of orthogroups – a core proteome total of 748 orthogroups (first bar from left). All other subsequent permutations of intersects contain less than 72 orthogroups.

These results indicate that wCin2USA1 is highly similar to the wMel strain at the genomic level while encoding a distinct proteomic profile. Such differences could influence host interactions, reproductive manipulation efficiency, and adaptation dynamics.

### Core Metabolic Pathways of Wolbachia spp. Remain Intact in the wCin2USA1 Genome

Biosynthetic pathways of interest in the context of *Wolbachia*-host symbiosis were identified through subsequent visualization and manual annotation (Fig 7), revealing complete pathways for riboflavin, purines, pyrimidines, and haem biosynthesis. Most of these pathways were found to be present in all *Wolbachia* genomes. Moreover, all strains were found to contain a suite of metabolite transport and secretion systems including those for haem, zinc, iron (III), lipoproteins, and phospholipids.

**Fig 7.**
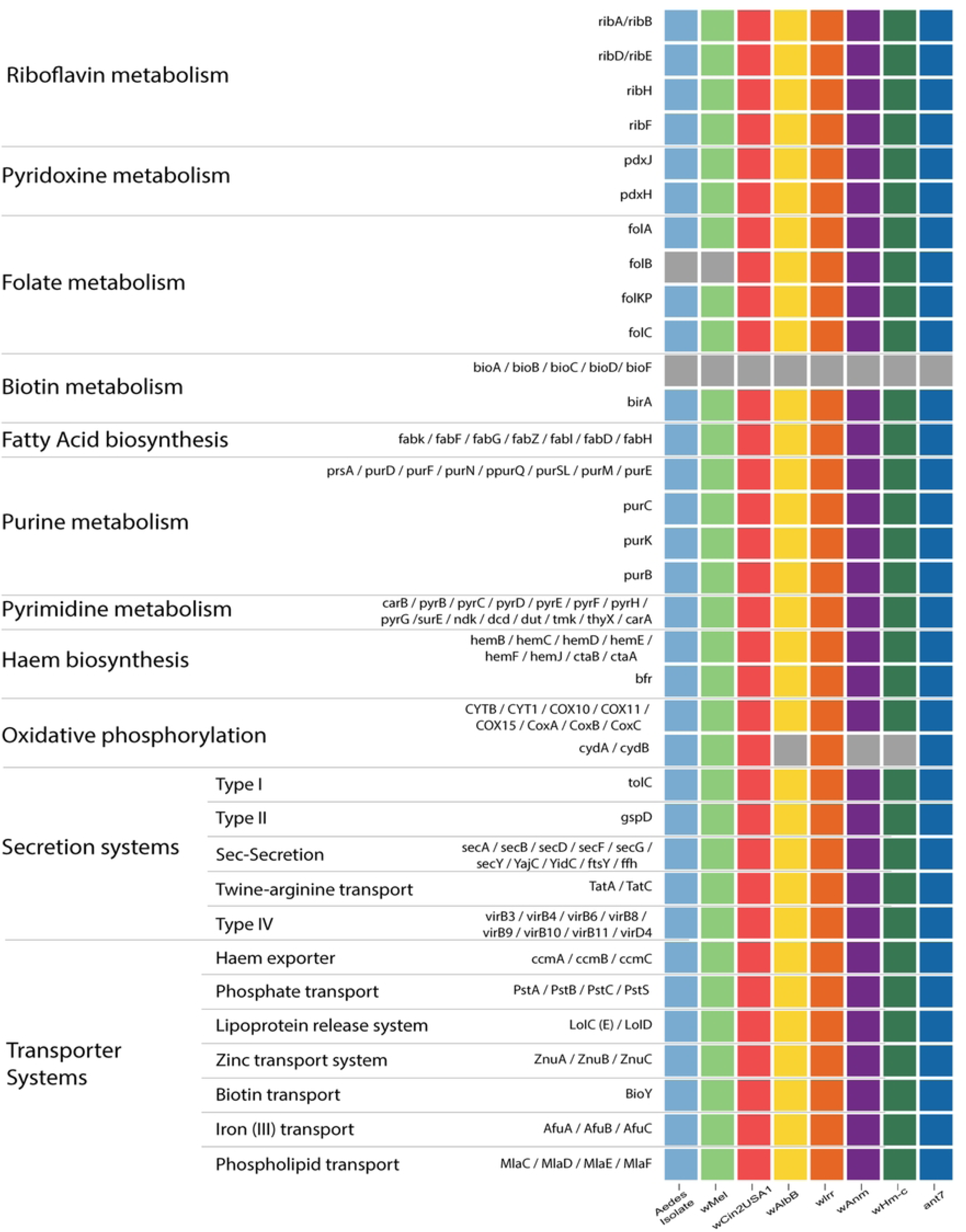
Color-wise representation of the presence–absence of various genes in metabolic and secretion/transport system pathways amongst the selection of *Wolbachia* genomes. The analyzed genomes are arrayed on the x-axis, with colors of the heatmap representing the various analyzed strains. The y-axis in turn represents different genes and metabolic pathways of interest to *Wolbachia* studies. Greys in the heatmap represent an absence of a gene within the respective genome.

All *Wolbachia* strains exhibit consistent metabolic and transport routes, supporting prior research findings. No biosynthetic pathways were uncovered that could suggest a new trait acquired in any of these strains. Relevant processes for *Wolbachia* include haem and nucleotide biosynthesis, as well as transport components like the T4SS for secreting protein effectors showcased to be present in all of the strains.

Although wAlbB, wAnm, and wHm-c strains lacked CydA and CydB genes, these genes were found in the remaining five strains. On the other hand, the *Aedes aegypti* isolate strain and wMel strain lack the folB gene, while the other six strains possess it. The wCin2USA1 strain, in addition to wlrr and ant7 strains, demonstrated the preservation of all genes associated with the pathways examined in this study, suggesting that wCin2USA1 possesses a wCin2USA1 possesses a more complete and conserved set of metabolic and functional pathways compared to the other strains, indicating greater genomic stability and potential functional versatility within the pathways analyzed in this study.

Additionally, the T4SSs and Sec-secretion systems were found to be maintained in all eight *Wolbachia* genomes. The T4SSs are known to play roles in infection and survival for a diverse range of symbiotic and pathogenic intracellular bacteria. It has been predicted that *Wolbachia* strains in the filarial nematode *Brugia malayi* utilize their T4SS to secrete protein effector molecules to avoid autophagy pathways and aid in actin cytoskeleton reformation, allowing intracellular mobility (Carpinone et al., 2018). Such processes may also be conserved within the *Wolbachia* strains analyzed in this study.

From the cluster of orthologous groups (COG) heatmap, the predominance of genes in COG category L (replication) within the core can be noticed (Fig 8). It is the most represented functional COG category in the *Wolbachia* genomes. The second most represented functional category contains genes involved in translation (J) followed by the S category that comprises of genes with unknown functions. Other significant COG categories that are distributed through all analyzed eight strains of *Wolbachia* are C, energy production and conversion; D, cell cycle control; E, amino acid transport and metabolism; F, nucleotide transport and metabolism and O, post-translational modification.

**Fig 8.**
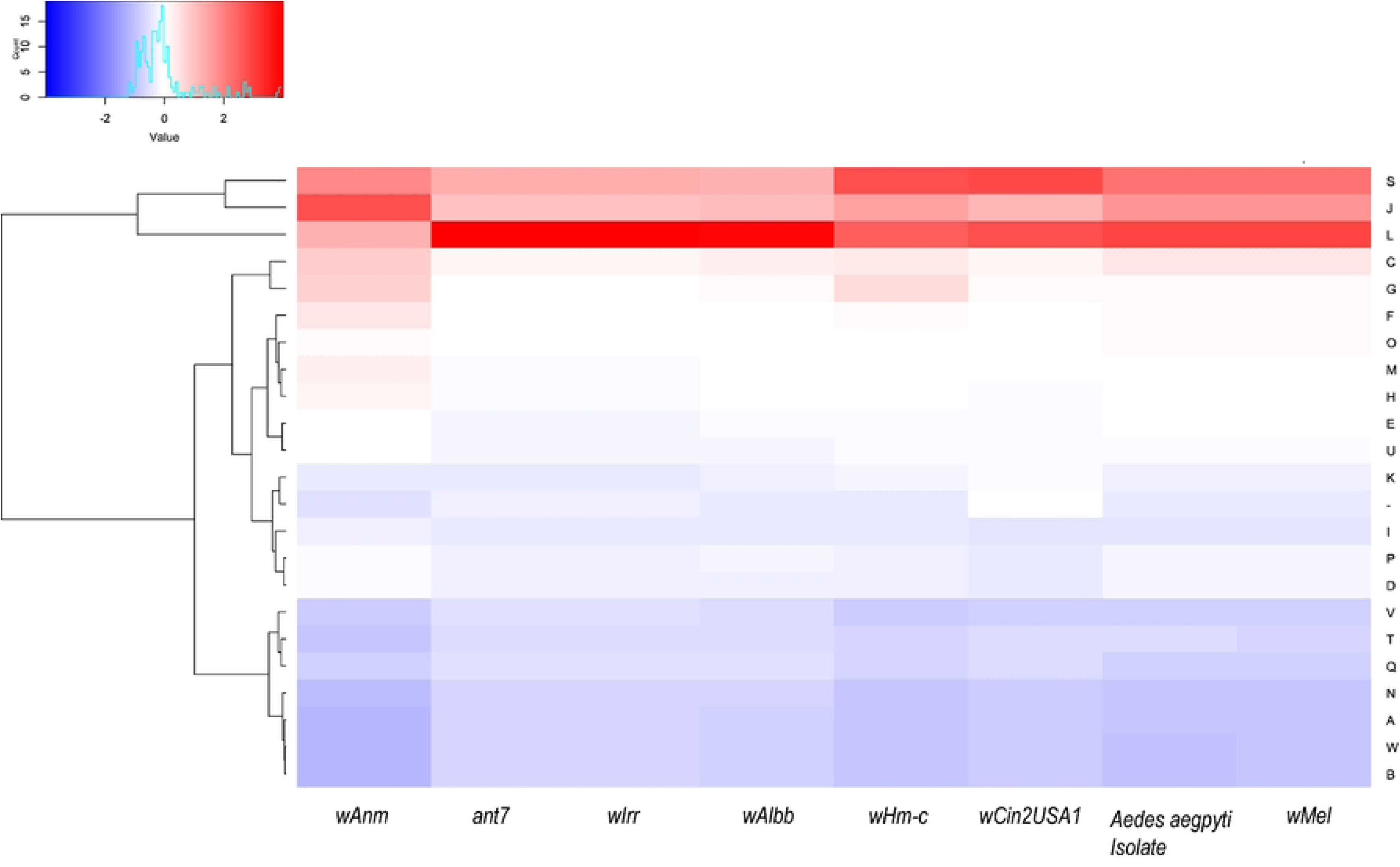
Heatmap comparison of the cluster of orthologous groups (COG) frequency profiles among *Wolbachia* strains. Abbreviations for functional categories are by alphabetic order: A, RNA processing and modification; B, Chromatic Structure and Dynamics; C, energy production and conversion; D, cell cycle control; E, amino acid transport and metabolism; F, nucleotide transport and metabolism; G, carbohydrate transport and metabolism; H, coenzyme transport and metabolism; I, lipid transport and metabolism; J, translation; K, transcription; L, replication, recombination, and repair; M, cell/wall membrane biogenesis; N, cell motility; O, post-translational modification, protein turnover, chaperones; P, inorganic ion transport and metabolism; Q, secondary metabolites biosynthesis, transport, and catabolism; R, general function prediction only; S, function unknown; T, signal transduction mechanism; U, intracellular trafficking and secretion; and V, defense mechanism

### Cif Gene Characterization revealed multiple Cif Genes present in the wCin2USA1 strain

Our genomic analysis of the *Wolbachia* strain wCin2USA1, previously uncharacterized and unreported in this context, identified two distinct syntennic cifA–cifB gene pairs, each affiliated with separate monophyletic clades within the cif gene family (Fig 9). This suggests that wCin2USA1 may harbor functionally and evolutionarily divergent cif operons, possibly acquired through separate horizontal transfer events. Notably, pair-1 of wCin2USA1 shows strong sequence similarity to the cif genes found in the *Aedes aegypti* isolate and the wMel strain, both of which belong to the Type-I monophyletic clade. In contrast, pair-2 of wCin2USA1 exhibits high homology to the cif genes of wAlbB, which fall within the Type-IV monophyletic clade ( Table 2).

**Fig 9.**
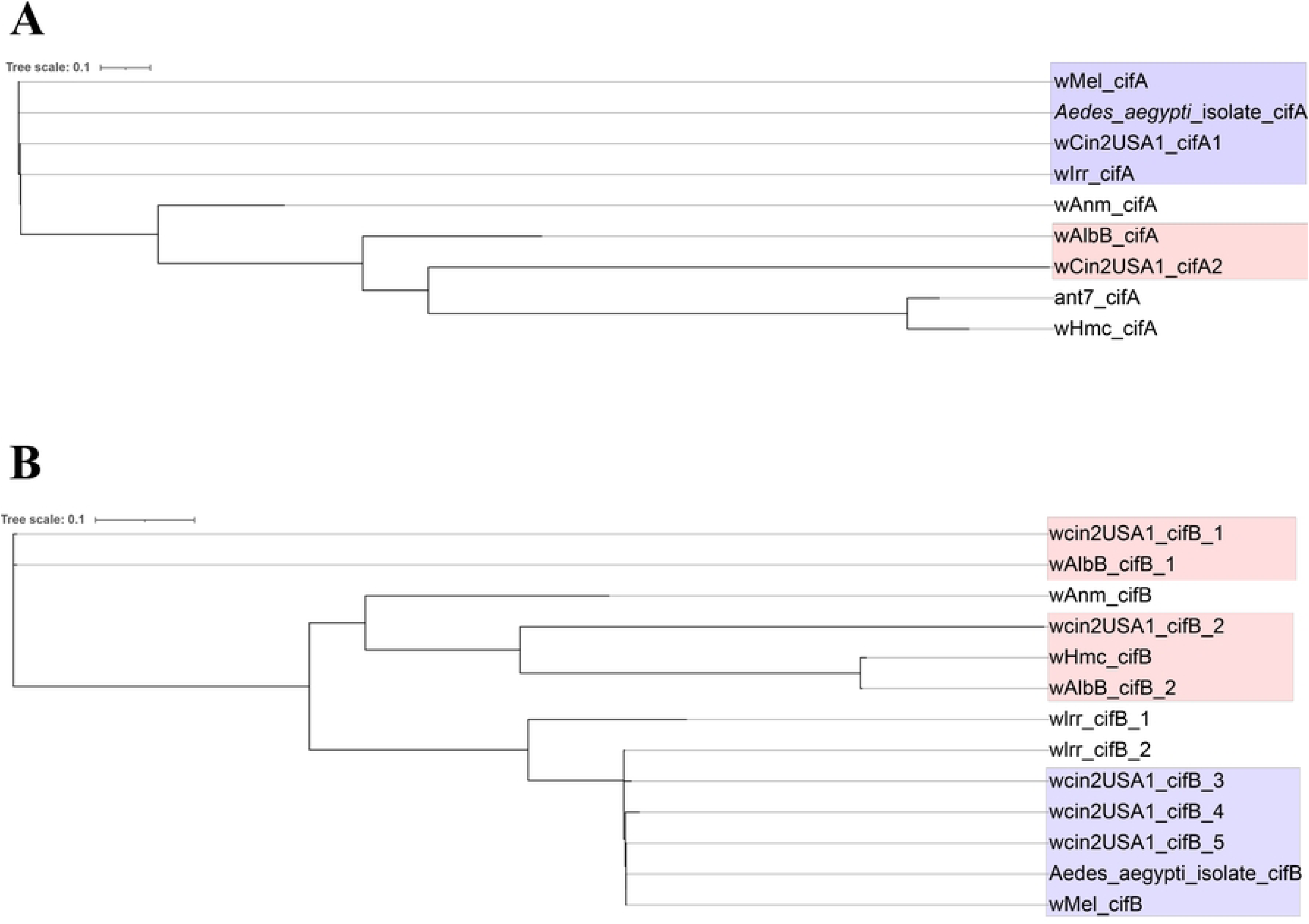
Phylogenetic analysis of cifA and cifB homologs in *Wolbachia* strain wCin2USA1 and related taxa. (A) Maximum likelihood phylogenetic tree of cifA genes, showing the relationship of wCin2USA1_cifA1 and wCin2USA1_cifA2 with cifA homologs from other *Wolbachia* strains. wCin2USA1_cifA1 clusters with the wMel and wLrr group (Type I), while wCin2USA1_cifA2 is more closely related to wAlbB_cifA (Type III), indicating the presence of two distinct monophyletic lineages of the wCin2USA1 cifA genes. (B) Phylogenetic tree constructed using cifB gene sequences highlights the presence of five cifB homologs in wCin2USA1, grouped into at least three distinct monophyletic clades. wCin2USA1_cifB_1 and wCin2USA1_cifB_2 cluster with wAlbB CifB homologs (Type III), while wCin2USA1_cifB_3,4 and 5 genes are more closely related to wMel and *Aedes aegypti* isolate cifB (Type I).

**Table 2.** Cif Gene Homologues of eight *Wolbachia* strains and their corresponding monophyletic types.

| Cif Gene Homologues | Cif Gene Monophyletic Type |
| --- | --- |
| *Aedes aegypti* Isolate Cif Gene | Type I |
| wMel Cif Gene | Type I |
| wCin2USA1 Cif Gene Pair 1<br>(wcin2USA1_cifA1 and wcin2USA1_cifB4) | Type I |
| wCin2USA1 Cif Gene Pair 2<br>(wcin2USA1_cifA2 and wcin2USA1_cifB1) | Type III |
| wIrr Cif Gene | Type I |
| wAlbB Cif Gene Pair 1 | Type IV |
| wAlbB Cif Gene Pair 2 | Type III |
| wHm-c Cif Gene | Type IV |
| wAnm Cif Gene | Type V |

*Wolbachia* genomes often encode multiple cifA–cifB gene pairs, a configuration thought to underlie complex patterns of bidirectional cytoplasmic incompatibility (Bonneau et al., 2018). The presence of two evolutionarily distinct cif operons in wCin2USA1 suggests that this strain may possess a hybrid CI-inducing potential, theoretically capable of mimicking the mechanisms of both wMel and wAlbB. Such duality could provide wCin2USA1 with increased flexibility and efficacy in manipulating host reproduction and blocking pathogen transmission (Beckmann et al., 2017).

Phylogenetic inference indicates that these gene pairs have diverged significantly from each other, aligning with distinct clades characterized in previous studies, such as those described by Martinez et al. (2021). This diversity could point to multiple functional mechanisms or evolutionary strategies for host manipulation.

In addition to the paired genes, three cifB genes were identified without an associated cifA partner in their genomic vicinity. The absence of adjacent cifA genes raises several possible interpretations. One explanation is that these cifB-only loci may represent degenerate or remnant operons, where the cifA gene has been lost due to decay or deletion, while the cifB component remains intact. Alternatively, these unpaired cifB genes might function independently or in conjunction with non-canonical partners. Some studies have proposed that cifB can retain partial or modified functionality even in the absence of cifA, particularly in strains where cytoplasmic incompatibility (CI) is weak or absent (Beckmann et al., 2017).

### Multiple Intact Prophage Regions Present in wCin2USA1

Analysis revealed that *wcin2USA1* maintains multiple intact and questionable prophage regions, including well-defined modules encoding structural phage proteins such as tail, head, portal, and terminase proteins, in contrast to several other *Wolbachia* strains where such regions are either degenerated or entirely absent. In total, seven prophage regions were identified, of which all seven appear to be intact based on the presence of hallmark phage genes (Fig 10). This pattern is consistent with prior observations in several *Wolbachia* strains that exhibit cytoplasmic incompatibility (CI), where prophage insertions remain partially or fully functional and retain conserved gene synteny (Lindsey et al., 2018).

**Fig 10.**
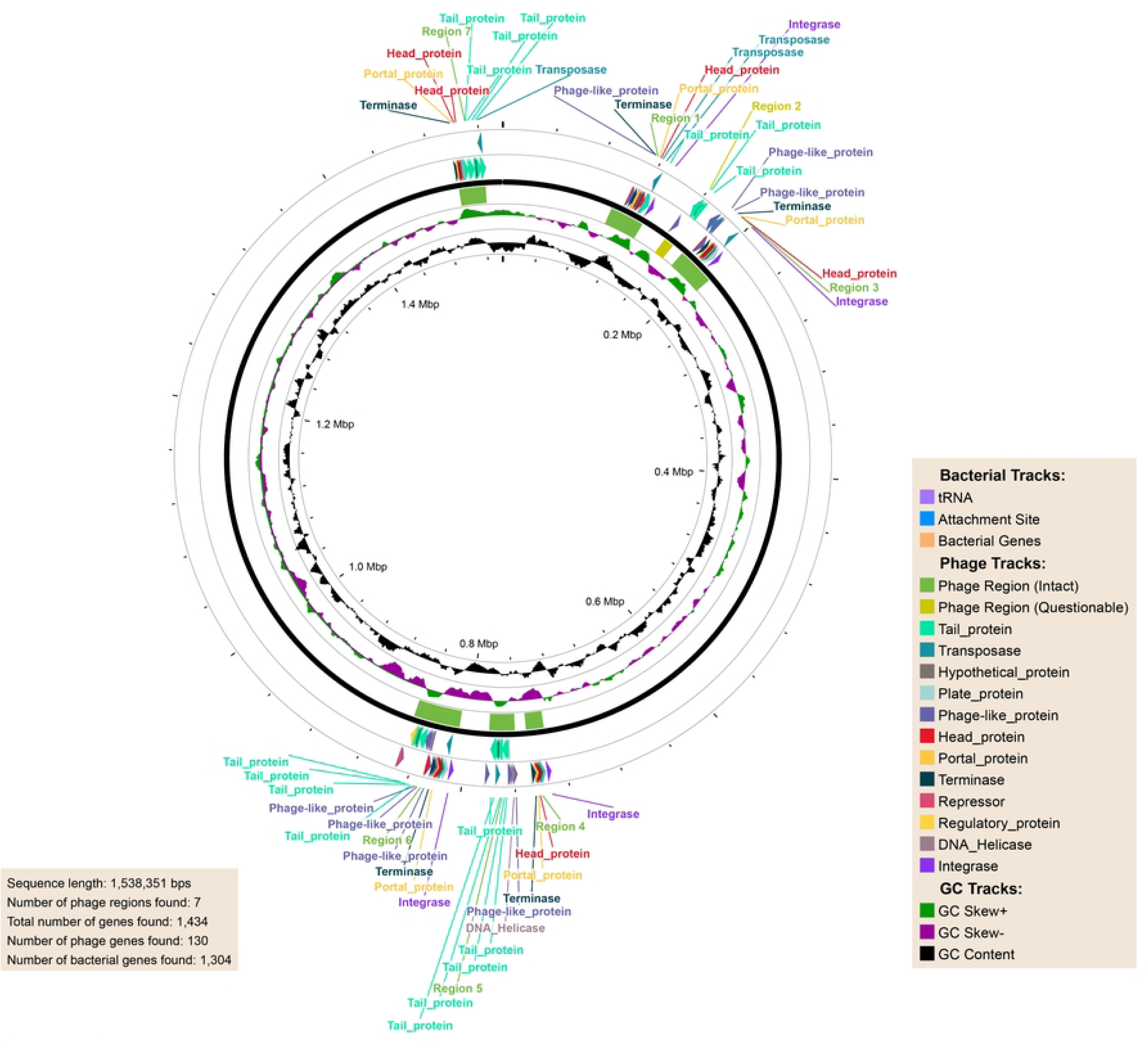
Circular genome map of wcin2USA1 highlighting prophage regions and associated phage genes.

This circular map represents the complete genome of the wcin2USA1 *Wolbachia* strain (1,538,351 bp), annotated for prophage regions and associated gene features. The outermost track marks the genomic coordinates in megabase pairs (Mbp). Intact prophage regions are shown in green and questionable regions in grey. Genes associated with phage elements are color-coded based on function: tail proteins (cyan), head proteins (red), portal proteins (orange), terminase (blue), phage-like proteins (purple), integrases (magenta), transposases (yellow), and other functional modules such as repressors and DNA helicases.

The presence of multiple integrases, transposases, and recombination-related proteins in *wcin2USA1* further supports the notion of ongoing or recent prophage activity, possibly enabling horizontal gene transfer or gene rearrangements within the *Wolbachia* genome (Fortier & Sekulovic, 2013). This stands in contrast to strains such as *wAnM*, where prophage regions are completely absent, a feature previously reported only in nematode-infecting *Wolbachia* (Quek et al., 2022), (Foster et al., 2005).

### Cytoplasmic Incompatibility Genes of wCin2USA1 Are Independent of Prophage Regions

None of the identified cif genes neither the paired nor unpaired are located within predicted prophage regions of the wCin2USA1 genome. This is notable given the well-documented association of cif genes with prophage WO in many *Wolbachia* strains (Kent & Bordenstein, 2010). The decoupling of cif genes from prophage contexts in wCin2USA1 may suggest either a genomic translocation event from their ancestral phage regions or long-term stabilization and domestication of these genes within the *Wolbachia* chromosome (LePage et al., 2017).

Such observations align with emerging evidence that cif genes, once integrated via phage-mediated transfer, can persist and evolve independently of prophage elements. This separation may allow *Wolbachia* to retain host-manipulative capabilities even in the absence of active prophage machinery, potentially representing an adaptive benefit (Shropshire et al., 2018).

The presence of multiple and divergent cif operons independent of prophage regions, alongside solitary cifB genes, underscores the genetic complexity of reproductive manipulation in wCin2USA1. It is plausible that different cif loci contribute to distinct CI phenotypes, or that some are vestigial while others remain functionally active (Martinez et al., 2021). Functional studies would be necessary to assess expression levels, protein interactions, and phenotypic outcomes in host insects.

## Conclusions

Our findings support the potential of *Wolbachia* strain wCin2USA1 as a next-generation biocontrol agent for *Aedes aegypti*. Its unique genomic architecture, particularly the presence of distinct *cif* gene pairs, suggests enhanced cytoplasmic incompatibility and virus-blocking capabilities, even under conditions where current strains might fail. This makes wCin2USA1 a promising candidate for transinfection into mosquito populations, particularly in regions experiencing a decline in the effectiveness of existing *Wolbachia* strains. Future work should experimentally validate its efficacy and fitness *in vivo* to confirm its potential as a more resilient and sustainable solution for dengue control.

